# Restricting CAR T Cell Trafficking Expands Targetable Antigen Space

**DOI:** 10.1101/2024.02.08.579002

**Authors:** Erin A. Morales, Kenneth A. Dietze, Jillian M. Baker, Alexander Wang, Stephanie V. Avila, Fiorella Iglesias, Sabarinath V. Radhakrishnan, Erica Vander Mause, Michael L. Olson, Wenxiang Sun, Ethan Rosati, Sadie L. Chidester, Thierry Iraguha, Xiaoxuan Fan, Djordje Atanackovic, Tim Luetkens

## Abstract

Chimeric antigen receptor (CAR) T cells are an effective treatment for some blood cancers. However, the lack of tumor-specific surface antigens limits their wider use. We identified a set of surface antigens that are limited in their expression to cancer and the central nervous system (CNS). We developed CAR T cells against one of these antigens, LINGO1, which is widely expressed in Ewing sarcoma (ES). To prevent CNS targeting, we engineered LINGO1 CAR T cells lacking integrin ⍺_4_ (A4^ko^), an adhesion molecule essential for migration across the blood-brain barrier. A4^ko^ LINGO1 CAR T cells were efficiently excluded from the CNS but retained efficacy against ES. We show that altering adhesion behavior expands the set of surface antigens targetable by CAR T cells.

**One sentence summary:** Altering integrin-mediated adhesion provides tumor selectivity to CAR T cells by preventing homing to defined normal tissues but retaining tumor trafficking and anti-tumor activity.

**GRAPHICAL ABSTRACT:** 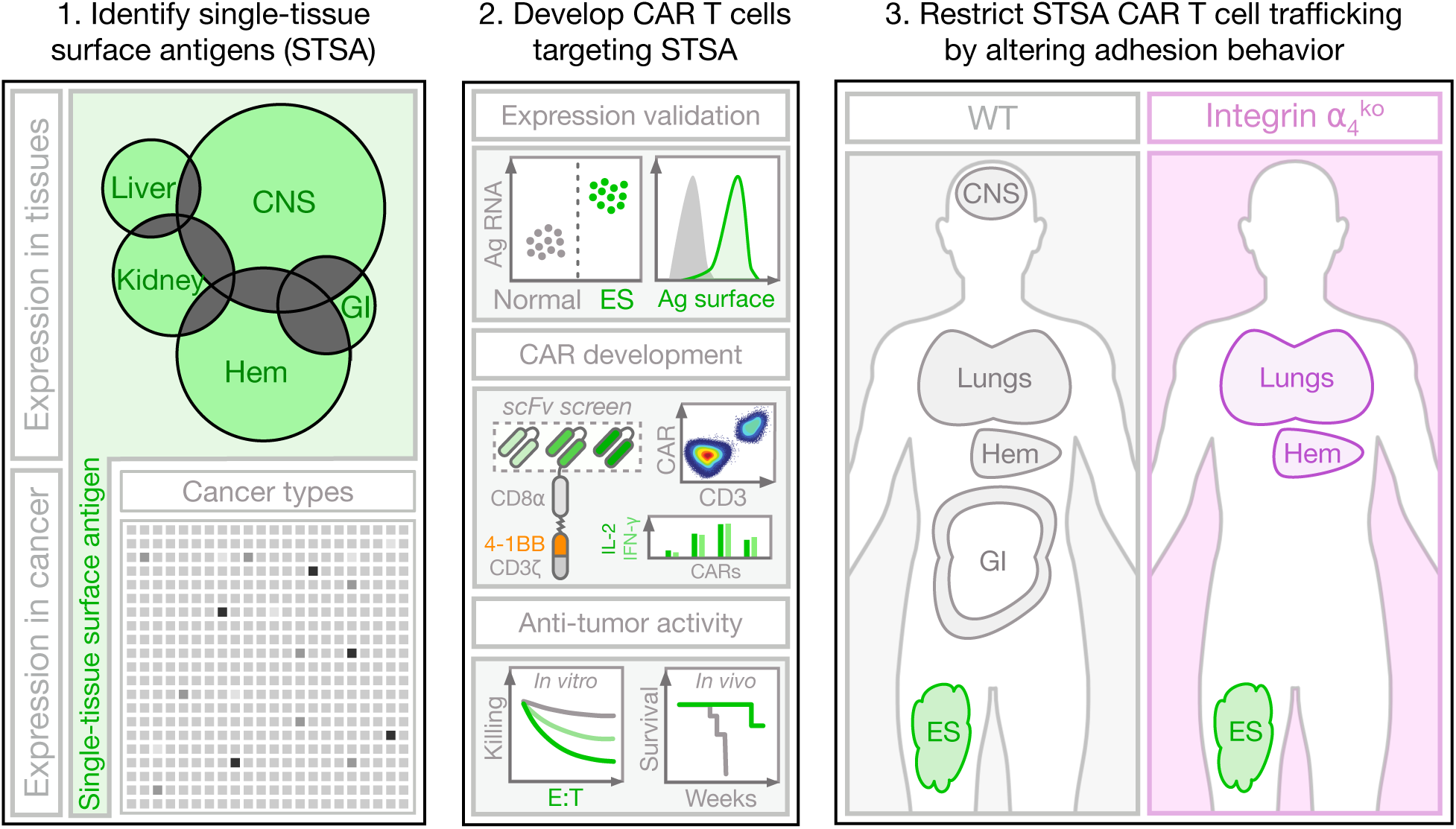

## INTRODUCTION

Chimeric antigen receptor (CAR) T cells are an effective treatment for cancer (*1–5*) and autoimmune diseases (*6*, *7*). Currently, six CAR T cell approaches targeting the antigens CD19 and BCMA are approved for the treatment of hematological cancers (*8*). However, a key challenge limiting the use of CAR T cells for other cancer types is the lack of surface antigens exclusively expressed on cancer cells. This lack of tumor specificity would result in the unintended targeting of normal tissues (on-target off-tumor toxicity) (*9*), preventing the exploration of otherwise promising tumor-associated surface antigens. Several approaches have been developed and tested preclinically to address this issue, such as preventing CAR T cell activation in the absence or presence of a second antigen (*10–12*), or by using strategies to induce CAR T cell expression only within defined tissues (*13*). These approaches rely on relatively homogeneous expression or lack of expression of these secondary antigens across the relevant target cells or tissues. An alternative strategy to limit on-target off-tumor toxicity could be to prevent CAR T cells from trafficking to normal tissues that express the target antigen. In the context of autoimmune diseases, altering T cell migration is a well-established and effective therapeutic strategy. For example, antibodies blocking T cell adhesion events to prevent their tissue-specific extravasation are an effective approach to reduce disease activity in multiple sclerosis (MS) (*14*) and Crohn’s disease (*15*, *16*). We hypothesize that altering T cell adhesion behavior could be an effective approach to provide tissue selectivity to engineered T cells and prevent the on-target off-tumor toxicity of CAR T cells, thereby expanding the targetable antigen space.

## RESULTS

### Single-tissue surface antigens are enriched in the CNS and expressed in non-CNS cancers

To determine whether restricting CAR T cell trafficking to reduce on-target off-tumor toxicity is feasible and likely to expand the set of targetable surface antigens, we first quantified the number of tumor-associated antigens that show restricted expression in normal tissues. Using publicly available consensus tissue RNA expression data (*17*) (Fig. 1A), we determined the number of antigens showing expression limited to a single normal tissue. We found that such single-tissue antigens are relatively common (*n* = 1,127), with the highest overall number of single-tissue antigens present in testis (Fig. 1B, Suppl. Table S1). When selecting single-tissue antigens that are predicted to be located on the cell surface, we found that such single-tissue surface antigens (STSAs) were most enriched in the central nervous system (CNS, Fig. 1B).

**Figure 1:**
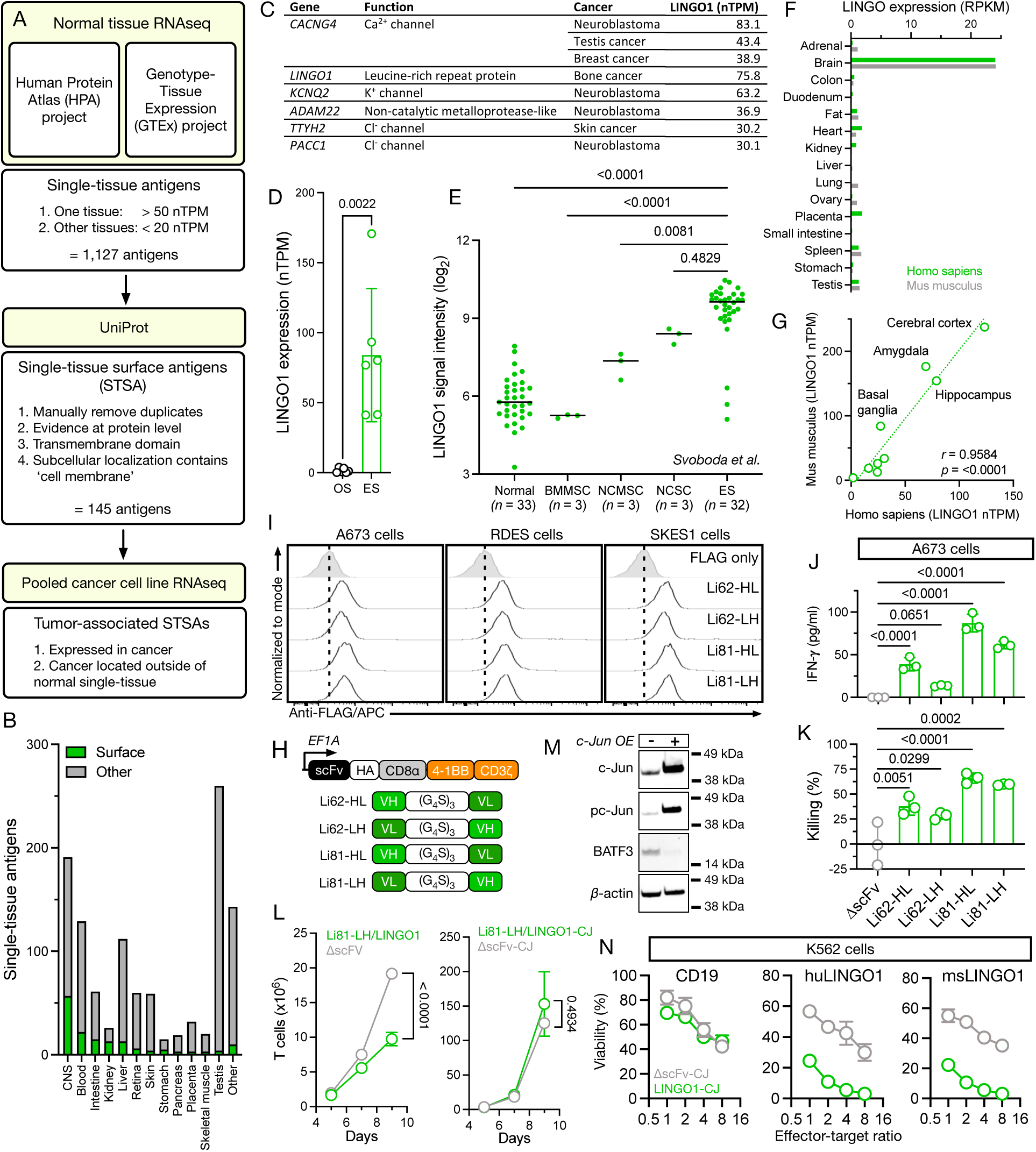
Tumor-associated single-tissue surface antigens are enriched in the CNS and can be targeted using CAR T cells. **(A)** Flowchart outlining process for the identification of single-tissue surface antigens using consensus normal tissue and cancer cell line RNA sequencing (RNAseq) data. **(B)** Frequency of single-tissue surface and non-surface antigens expressed in different normal tissues. **(C)** List of CNS-restricted STSAs with highest RNA expression levels in cancer. **(D)** RNA expression of LINGO1 in osteosarcoma (OS) and Ewing sarcoma (ES) cell lines as determined by RNAseq. Data represent mean ± S.D. from 6 cell lines per diagnosis. Statistical significance was determined by Mann-Whitney U test. **(E)** Expression of LINGO1 in primary human ES tumor biopsies, normal tissues, bone marrow mesenchymal stem cells (BMMSC), neural crest mesenchymal stem cells (NCMSC), and neural crest stem cells (NSCS) as determined by HuEx microarray (*20*). Data represent mean ± S.D. from indicated number of patients or tissues. Statistical significance was determined by one-way ANOVA. **(F)** LINGO1 expression in normal human and mouse tissues as determined by RNAseq. RPKM = reads per kilobase per million mapped reads. **(G)** LINGO1 expression in human and mouse brain regions as determined by RNAseq. **(H)** Schema of LINGO1 CAR and scFv constructs. **(I)** Surface expression of LINGO1 on ES cell lines using single-chain variable fragments derived from monoclonal LINGO1 antibody clones Li62 and Li81 as determined by flow cytometry. **(J)** IFN-γ secretion and **(K)** target cell killing by LINGO1 CAR T cell variants after 16-hour incubation with ES cell line A673 expressing firefly luciferase at an effector-target ratio of 1:1. Data represent mean ± S.D. from 3 technical replicates. Statistical significance was determined by one-way ANOVA. **(L)** Expansion of LINGO1 CAR T cells with and without overexpression of c-Jun during manufacturing. Data represent mean ± S.D. from 3 donors. Statistical significance was determined by two-sided Student’s t test. **(M)** Expression of transcription factors c-Jun and BATF3 in LINGO1 CAR T cells with and without overexpression of c-Jun as determined by western blot. **(N)** Killing of K562 cells engineered to express CD19, human LINGO1 (huLINGO1), or mouse LINGO1 (msLINGO1) following 16-hour co-culture with c-Jun (CJ)-expressing T cells expressing an Li81-based LINGO1 CAR or a control CAR without a binding domain (ΔscFV). Data represent mean ± S.D. from 3 technical replicates.

We next determined whether the identified STSAs show expression in human cancer (*18*), disregarding any STSAs for which tumors would be primarily located within an STSA’s respective normal tissue (e.g. expression of CNS-restricted STSAs in brain tumors). We found that CNS-restricted STSAs are commonly expressed in several non-CNS cancers (Suppl. Fig. S1A, Suppl. Table S2). By preventing CAR T cells from trafficking to the CNS, these antigens may represent a promising resource for the development of new CAR T cell approaches.

### The CNS-restricted STSA LINGO1 is expressed in Ewing sarcoma and can be targeted using LINGO1 CAR T cells

To determine if CNS-restricted STSAs can be targeted using CAR T cells, we first explored those STSAs with the highest expression in non-CNS tumors (Fig. 1C). Among these highly expressed antigens, we identified several ion channels, which can be difficult to target using antibody-based approaches due to their small extracellular footprint and high structural homology to each other (*19*). However, we also identified proteins with large unique extracellular domains that could represent promising targets for CAR T cell approaches, such as leucine-rich repeat and Immunoglobin-like domain-containing protein 1 (LINGO1), a type 1 transmembrane protein expressed in Ewing sarcoma (ES, Fig. 1D, Suppl. Fig. S1B). Analyzing LINGO1 expression data from primary human samples (*20*), we observed increased expression in candidate cells of ES origin with the highest expression in human ES tumors (Fig. 1E). Expression above normal tissues was observed in 29/32 (90.6%) ES tumors indicating that therapeutic targeting of LINGO1 may be possible in a majority of patients with ES. Gene expression within ES tumors, even of the *EWSR1-FLI1* fusion gene driving the malignant ES phenotype, is highly heterogeneous (*21*, *22*). However, among candidate antigens currently considered for CAR T cell-based treatments of ES (*23*, *24*), *LINGO1* showed the most uniform expression (Suppl. Fig. S1C) (*25*). In normal tissues, we observed conserved LINGO1 expression restricted to the CNS (Fig. 1F), specifically the cerebral cortex, amygdala, and hippocampus (Fig. 1G). These data validate LINGO1 as a CNS-restricted STSA and potential target for the treatment of ES using CAR T cells.

Despite the otherwise tumor-specific expression of LINGO1, no CAR T cell approaches targeting this antigen have been developed. Monoclonal antibodies targeting the extracellular domain of LINGO1 were described previously (*26*) and the LINGO1-specific antibody opicinumab has been tested clinically for the treatment of MS (*27*). To produce LINGO1-specific CAR T cells, we generated single-chain variable fragments (scFv) based on two of these existing antibodies (Fig. 1H), clones Li62 and Li81, as potential CAR binding domains. All four scFvs could be expressed at concentrations sufficient for immunofluorescence staining (Suppl. Fig. S1D) and showed binding to all 3 of the ES cell lines tested with only minimal differences between clones and variable heavy (VH)/variable light (VL) chain orientation (Fig. 1I). Positive surface staining of ES cells using anti-LINGO1 scFvs was validated using opicinumab (Suppl. Fig. S1E). We next generated 4-1BB-based second-generation CAR T cells (Fig. 1H) using the four scFvs as well as a CAR lacking a binding domain (ΔscFv) to serve as negative control. All constructs showed surface expression when introduced into primary human T cells (Suppl. Fig. S1F). When co-cultured with the ES cell line A673, LINGO1 CAR constructs showed varying levels of IFN-γ production (Fig. 1J) and ES tumor cell killing (Fig. 1K), with constructs based on clone Li81 showing superior activity. While Li81 in VH-VL orientation showed slightly higher IFN-γ production and tumor cell killing, we selected the VL-VH orientation for subsequent experiments due to its substantially higher surface expression.

During CAR T cell production, we observed reduced expansion of LINGO1 CAR T cells compared to ΔscFv CAR T cells (Fig. 1L), a hallmark of tonic signaling leading to early T cell exhaustion (*28*). It has previously been shown that overexpression of the transcription factor c-Jun can counteract tonic signaling in CAR T cells (*29*). We overexpressed c-Jun (Fig. 1M, Suppl. Fig. S1G) in LINGO1 (LINGO-CJ) and ΔscFv (ΔscFv-CJ) CAR T cells and observed reduced production of the exhaustion-associated transcription factor BATF3 (Fig. 1M) and rescued LINGO1 CAR T cell expansion (Fig. 1L), indicating ameliorated tonic signaling. To determine the antigen specificity and cytotoxic activity of LINGO1-CJ CAR T cells, we next generated K562 cells expressing firefly luciferase (Fluc) together with CD19, human LINGO1 (huLINGO1) (Suppl. Fig. S1H), or the highly conserved (Suppl. Fig. S1I) mouse LINGO1 (msLINGO1). LINGO1-CJ CAR T cells did not show killing of K562 cells expressing CD19 but efficiently lysed K562 cells expressing human or mouse LINGO1 (Fig. 1N). In summary, we show that LINGO1 expression is restricted to ES and the CNS and that CAR T cells are an effective approach for the specific targeting of LINGO1-expressing cells.

### CAR T cells targeting the CNS-restricted STSA LINGO1 efficiently eradicate Ewing sarcoma in vitro and in vivo

To date no CAR T cell approaches are approved for the treatment of ES. We hypothesize that LINGO1 CAR T cells could be an effective approach for the targeting of ES. Co-culturing human ES cell lines with LINGO-CJ CAR T cells (Fig. 2A, Suppl. Fig. S2A) results in efficient dose-dependent ES cell killing (Fig. 2B) and production of effector cytokines (Fig. 2C) even at low effector-target ratios. Primary ES localizes to the bone but can metastasize to the bone marrow and the lungs (*30*). To determine the long-term anti-tumor efficacy of LINGO1-CJ CAR T cells against ES tumors, we performed a metastatic ES mouse model with systemically injected human ES cell line A673 (Fig. 2D). Treatment of mice with ES tumors using a single dose of systemically injected LINGO1-CJ CAR T cells resulted in tumor eradication in all animals (Fig. 2E, Suppl. Fig. S2B) and significantly increased survival (Fig. 2F).

**Figure 2:**
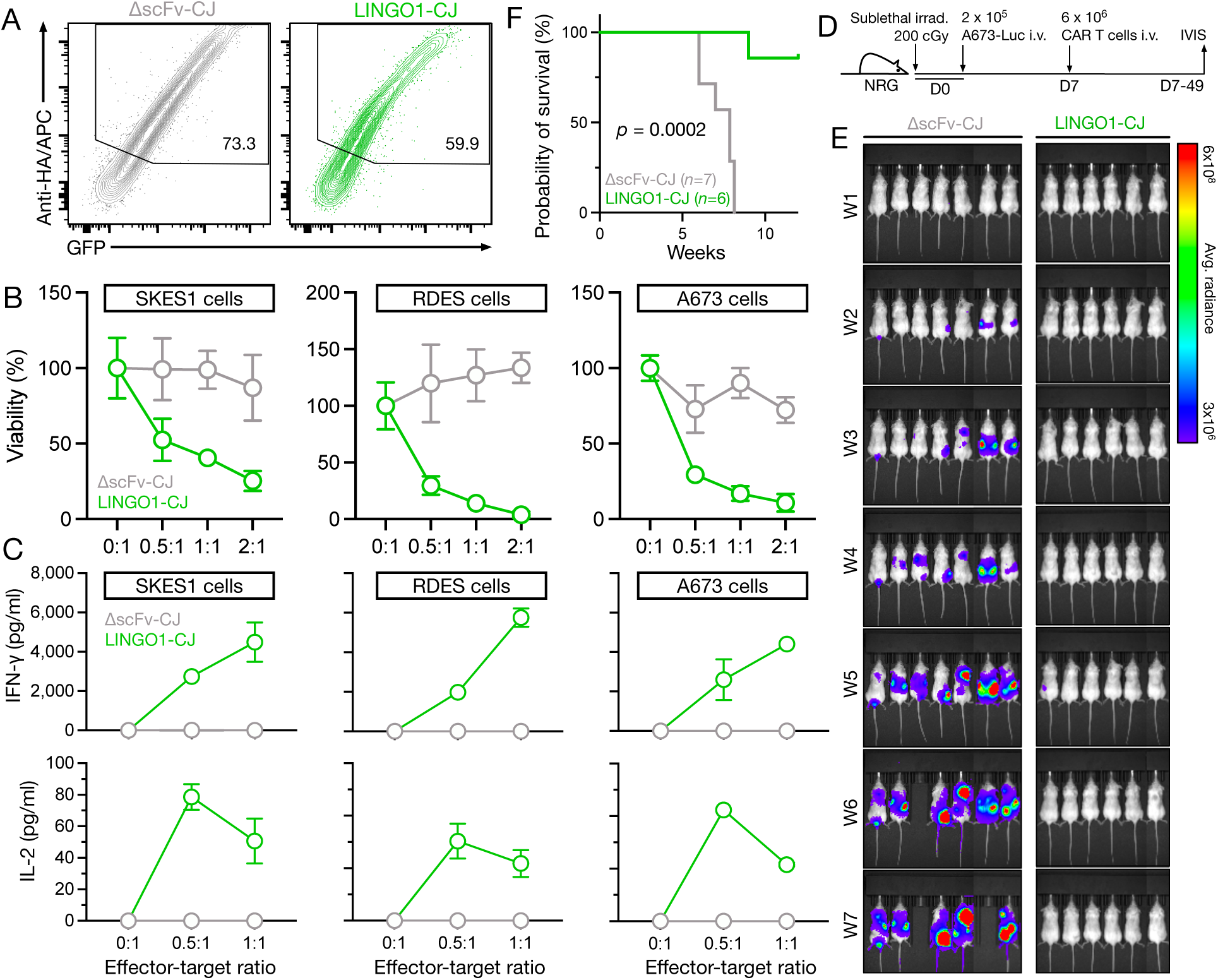
LINGO1 CAR T cells efficiently eradicate ES *in vitro* and *in vivo*. **(A)** CAR surface and GFP reporter expression in primary human T cells at the end of gammaretroviral manufacturing. Data are representative of 4 independent CAR T cell productions. **(B)** Killing of 3 ES cell lines after 16-hour co-culture with T cells expressing LINGO1 CAR or ΔscFv CAR at the indicated effector-target ratios as determined by luciferase-based cytotoxicity assay. Data represent mean ± S.D. from 3 technical replicates. **(C)** Cytokine concentrations in co-culture supernatants after 16-hour as determined by ELISA. Data represent mean ± S.D. from 3 technical replicates. **(D)** Schema of *in vivo* experiment to determine efficacy of LINGO1 CAR T cells. **(E)** Tumor burden in mice bearing systemic A673 tumors and treated with LINGO1 CAR T cells or ΔscFv control CAR T cells as determined by *in vivo* imaging system (IVIS). **(F)** Survival of ES tumor bearing mice treated with LINGO1 CAR T cells. Statistical significance was determined by log-rank test.

### Integrin ⍺_4_ knockout in human T cells is feasible and abrogates adhesion to VCAM-1

The CNS is considered immune-privileged mainly due to its separation from components of the immune system by the blood-brain barrier (*31*). While trafficking by T cells and other immune cells to the CNS is tightly regulated, it is a well-established process that is essential to clearing infections within the CNS as well as the pathophysiology of different neurodegenerative autoimmune diseases (*32*). To initiate crossing of the blood-brain barrier, T cells rely on a single type of adhesion event for capture and sustained cell arrest: the binding of the activated integrin ⍺_4_β_1_ heterodimer on T cells to *VCAM-1* on brain endothelial cells (*33*) (Fig. 3A). Therapeutic inhibition of this process by the integrin ⍺_4_-specific monoclonal antibody natalizumab in patients with MS is highly effective at preventing T cell trafficking to the CNS and reducing disease activity (*14*). Importantly, the frequency of malignancies is not increased in these patients (*34*) suggesting that this adhesion event is not essential for the trafficking of T cells to tumors. It has, however, been shown that treatment is associated with a rare viral infection that can result in potentially fatal progressive multifocal leukencephalopathy (PML) (*35*, *36*), likely due to a global reduction in T cell surveillance of the CNS (*37*). To avoid this side effect, we explored targeting the ⍺_4_β_1_/VCAM-1 adhesion event genetically to alter the trafficking of CAR T cells only – allowing for non-engineered T cells to continue regular CNS immune surveillance.

**Figure 3:**
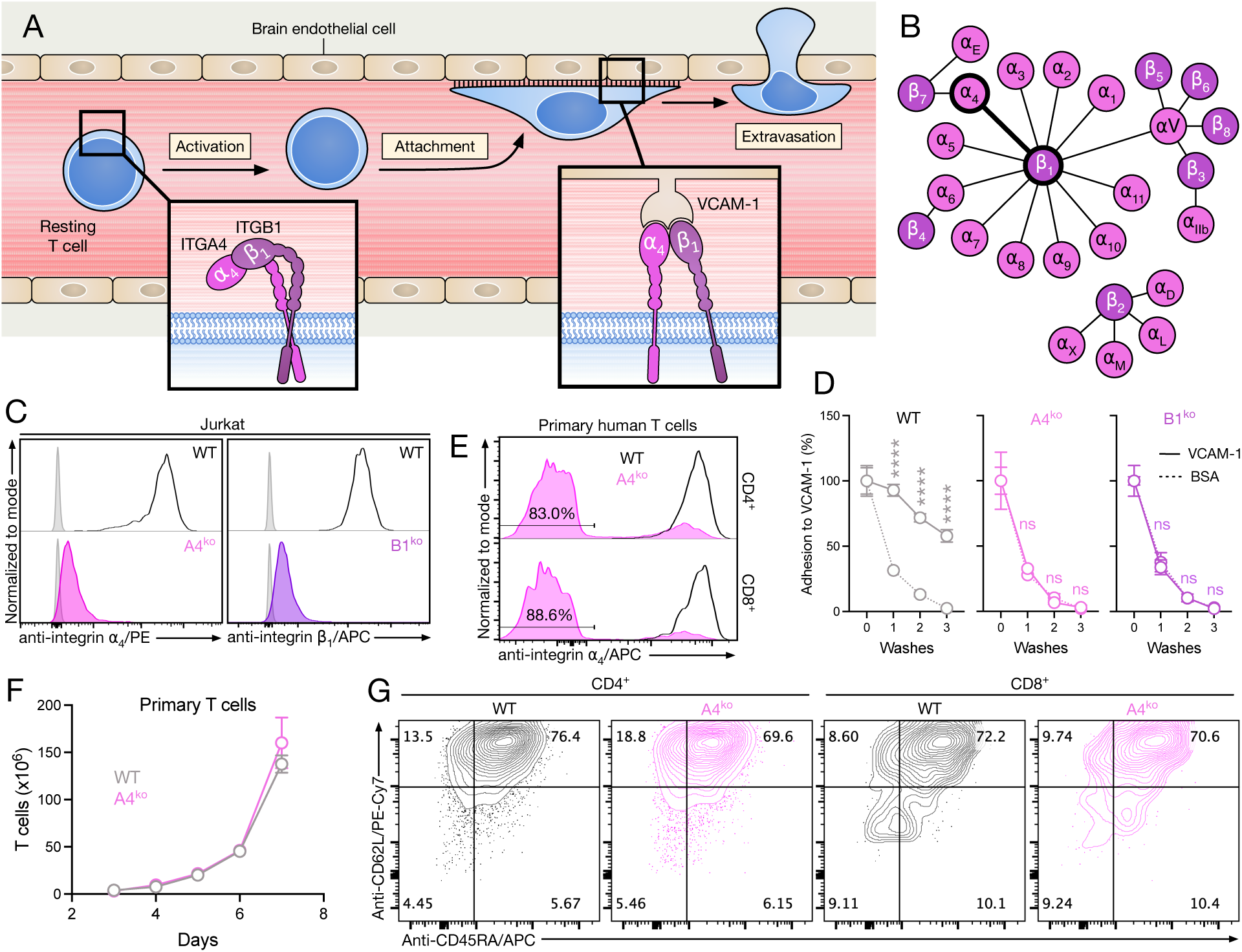
*Integrin ⍺-4* knockout in human T cells prevents adhesion to VCAM-1 and does not alter T cell expansion or phenotype. **(A)** Schema of integrin ⍺_4_-mediated T cell attachment and extravasation. **(B)** Schema of integrins and their respective heterodimerization partners. **(C)** Knockout efficiency of integrin ⍺_4_ and β_1_ in Jurkat cells transduced with a lentiviral construct expressing Cas9 and non-specific (WT), or integrin ⍺_4_ or integrin β_1_-targeting gRNA constructs as determined by flow cytometry after cell sorting. Unstained control in grey. **(D)** Adhesion of Jurkat-Fluc cells to recombinant VCAM-1 following integrin ⍺_4_ or integrin β_1_ knockout as determined by luminescence. Data indicate mean ± S.D. from 2 technical replicates. Statistical significance was determined by two-way ANOVA. **(E)** Integrin ⍺_4_ knockout efficiency in primary human T cells after electroporation with CRISPR/Cas9 RNP as determined by flow cytometry. **(F)** Expansion of stimulated T cells following integrin ⍺*_4_* knockout. Data represent mean ± S.D. from 3 different donors. **(G)** T cell phenotype 10 days after integrin ⍺_4_ knockout as determined by flow cytometry.

Integrins are an abundant class of proteins that are essential for cell adhesion (*38*). Some integrins can pair with multiple corresponding integrins (Fig. 3B) with each heterodimer conferring a specific adhesion behavior. Integrins ⍺_4_ and β_1_ are encoded by the *ITGA4* and *ITGB1* genes, respectively. Comparing multiple guide RNAs (Suppl. Fig. S3A), we found that CRISPR/Cas9-mediated functional knockout of *ITGA4* or *ITGB1* in Jurkat cells was feasible and efficient (Fig. 3C, Suppl. Fig. S3B). Knockout of either component of the integrin ⍺_4_β_1_ heterodimer abrogated binding of T cells to recombinant VCAM-1 (Fig. 3D). Due to the lower number of heterodimer combinations involving integrin ⍺_4_ and the established clinical safety of targeting integrin ⍺_4_, we selected integrin ⍺_4_ knockout (A4^ko^) to alter CNS-trafficking of CAR T cells. We found that CRISPR ribonucleoprotein (RNP)-mediated A4^ko^ was efficient in activated primary human T cells (Fig. 3E) and did not affect T cell expansion (Fig. 3F) or phenotype (Fig. 3G).

### A4^ko^ LINGO1 CAR T cells can be manufactured efficiently and retain anti-tumor activity against ES

While the primary function of integrins is to mediate cell-cell adhesion, integrins also contribute to signaling, cell motility, and proliferation (*39*, *40*). To determine if A4^ko^ alters essential CAR T cell functions, we developed a scalable process for the generation of A4^ko^ LINGO1 CAR T cells (Fig. 4A). The resulting product shows comparable CAR expression between WT and A4^ko^ CAR T cells and an average of 89% integrin ⍺_4_^neg^ T cells (Fig. 4B, Suppl. Fig. S4A). A4^ko^ did not significantly alter the surface expression of phenotypic, activation, or exhaustion markers at the end of CAR T cell production (Fig. 4C, Suppl. Fig. S4B) or the expansion of CAR T cells (Fig. 4D).

**Figure 4:**
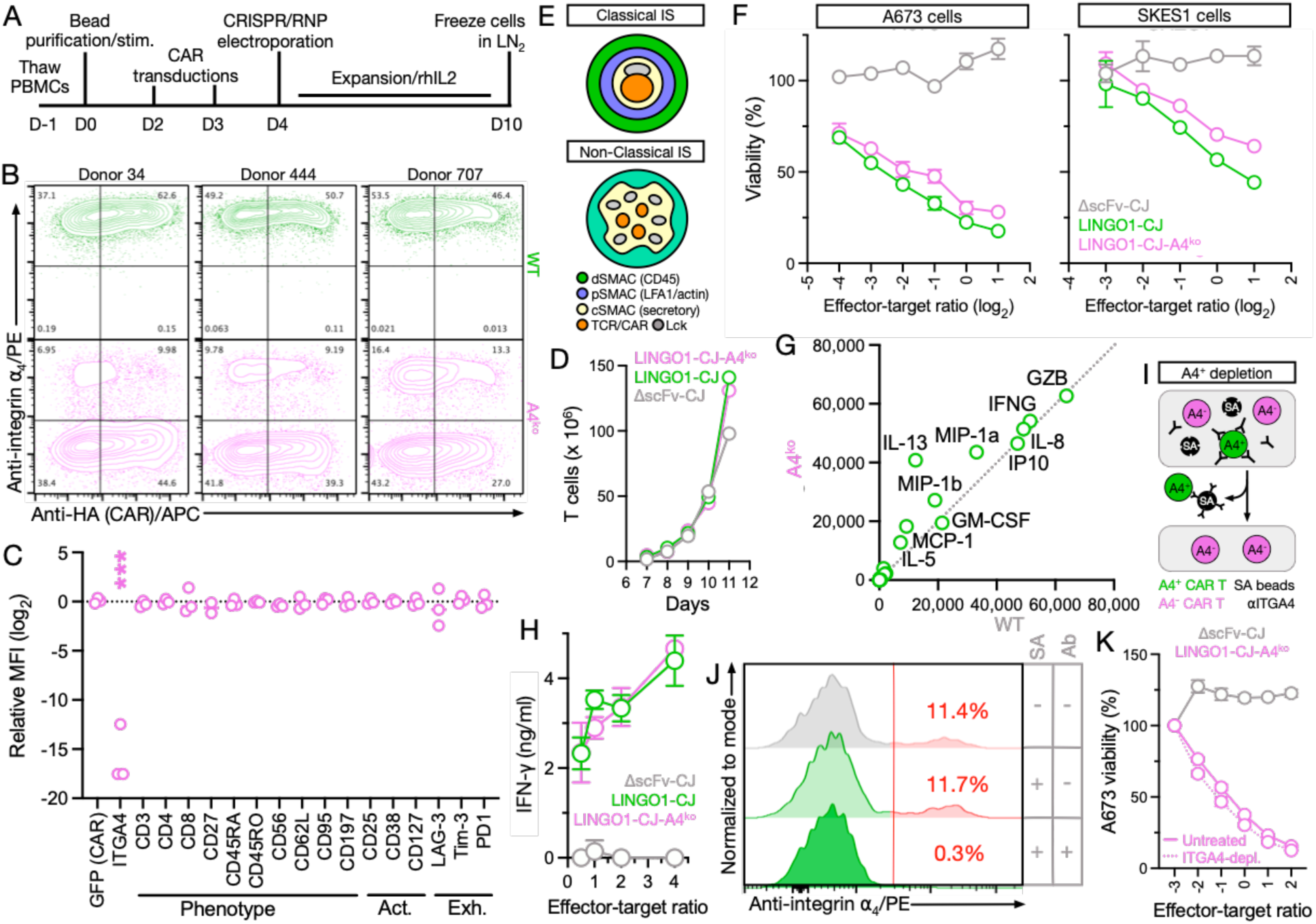
A4^ko^ LINGO1 CAR T cells can be manufactured efficiently and maintain anti-tumor activity against ES. **(A)** Schema of A4^ko^ LINGO1 CAR T cell manufacturing workflow. **(B)** Integrin ⍺_4_ knockout and CAR transduction efficiency in 3 donors as determined by flow cytometry. **(C)** Expression of T cell markers in WT and A4^ko^ LINGO1 CAR T cells from 3 healthy donors as determined by spectral flow cytometry. Data were normalized to the mean of the WT condition. Statistical significance of differences between WT and A4^ko^ LINGO1 CAR T cells was determined by two-way ANOVA. **(D)** Expansion of WT or A4^ko^ LINGO1 CAR T cells as determined by automated cell counting. **(E)** Schematic showing location of distal (d), peripheral (p), and central (c) supramolecular activation complex (SMAC) in classical and non-classical/CAR immune synapses (IS). **(F)** Killing of 2 ES cell lines after 16-hour co-culture with T cells expressing LINGO1 CAR or ΔscFv CAR at the indicated effector-target ratios as determined by luminescence cytotoxicity assay. Data represent mean ± S.D. from 3 technical replicates. **(G)** Cytokine concentrations in supernatants of WT or A4^ko^ LINGO1 CAR T cells co-cultured with A673 cells after 16 hours as determined by CodePlex assay and **(H)** ELISA. **(I)** Schema of streptavidin (SA)-bead-based approach for the depletion of remaining integrin ⍺ ^pos^ LINGO1 CAR T cells. **(J)** Quantification of *integrin ⍺-4*^pos^ LINGO1 CAR T cell following bead-based depletion as determined by flow cytometry. **(K)** Killing of A673 cells with or without depletion of integrin ⍺ ^pos^ LINGO1 CAR T cells as determined by luminescence-based cytotoxicity assay. Data represent mean ± S.D. from 3 technical replicates.

When T cells encounter target cells, they form immune synapses (IS) to coordinate molecular recognition, intracellular signaling, cytoskeletal rearrangement, and granule secretion resulting in target cell killing (*41*). While the formation of classical immune synapses in conventional T cells is well characterized, the non-classical CAR T cell synapse remains relatively poorly understood (Fig. 4E). A key difference between the classical and non-classical IS is the relative lack of contribution of the adhesion molecule *LFA-1* to CAR T cell-mediated killing (*42*). It is unknown if other adhesion molecules, such as integrin ⍺_4_, contribute to non-classical IS formation either in addition to or instead of *LFA-1*. We therefore determined if LINGO1 CAR T cells would be able to maintain their anti-tumor activity in the presence of integrin ⍺_4_ knockout. We found that A4^ko^ LINGO1 CAR T cells showed killing of ES cell lines (Fig. 4F) and a cytokine secretion profile (Fig. 4G/H) comparable to WT LINGO1 CAR T cells. To rule out the possibility that the remaining integrin ⍺_4_^pos^ LINGO1 CAR T cells were responsible for the observed effects, we established a process (Fig. 4I) for the efficient depletion of remaining integrin ⍺_4_^pos^ T cells from the final CAR T cell product (Fig. 4J, Suppl. Fig. S4C). We found that integrin ⍺_4_-depleted A4^ko^ LINGO1 CAR T cells exhibited a comparable amount of ES killing to non-depleted A4^ko^ LINGO1 CAR T cells, confirming that the observed effects were not due to remaining integrin ⍺_4_^pos^ CAR T cells and that A4^ko^ does not alter the in vitro anti-tumor activity of LINGO1 CAR T cells (Fig. 4K). Overall, A4^ko^ LINGO1 CAR T cells maintain expansion, phenotype, *in vitro* anti-tumor activity, and cytokine secretion of WT LINGO1 CAR T cells.

### Integrin ⍺_4_ knockout prevents trafficking of CAR T cells to the CNS but not to ES tumors

We hypothesize that integrin ⍺_4_ knockout enables the selective targeting of tumors expressing CNS-restricted antigens using CAR T cells by preventing trafficking to the CNS. To determine the effect of A4^ko^ on CAR T cell trafficking, we performed a series of *in vivo* experiments. Integrins ⍺_4_ and β_1_ as well as VCAM-1 are highly conserved between humans and mice (Suppl. Fig. S5A) and human integrin ⍺_4_β_1_ has been shown to bind efficiently to mouse VCAM-1 (43). Mouse models have been used previously to investigate human T cell trafficking generally recapitulating tissue homing observed in humans (*44–46*). Using human A4^ko^ LINGO1 CAR T cells engineered to co-express Fluc, we first determined CAR T cell trafficking to different organs in mice by IVIS (Fig. 5A). We found that A4^ko^ did not alter CAR T cell trafficking to the lungs and spleen but significantly reduced trafficking to the intestine and stomach (Fig. 5B/C), likely by disrupting formation of integrin ⍺_4_β_7_ which is necessary for trafficking to the gastrointestinal tract (*47*, *48*). In this short-term model, we were not able to detect either WT or A4^ko^ LINGO1 CAR T cells in the brains of animals by *in vivo* imaging system (IVIS) or flow cytometry (Suppl. Fig. S5B/C). Adhesion of integrin ⍺_4_β_1_ to VCAM-1 requires prior activation of the integrin heterodimer to bind VCAM-1 with high affinity (*49*). While resting primary T cells mainly express inactive low-affinity integrin ⍺_4_β_1_, the human T cell line Jurkat expresses a constitutively active form of integrin ⍺_4_β_1_ (*50*) (Fig. 5D). We hypothesized that Jurkat cells may be a more sensitive model to quantify CNS trafficking of human A4^ko^ T cells. Following systemic injection of WT or A4^ko^ Jurkat-Fluc cells, we performed IVIS imaging to analyze T cell trafficking to the CNS (Fig. 5E). A4^ko^ appeared to lead to an overall reduction in IVIS signal in the head of the animals (Fig. 5F/G) and analyzing explanted brains from the same animals, we observed significantly lower IVIS signal in the brains of animals injected with A4^ko^ Jurkat cells (Fig. 5H). Quantification of T cells in these brains by flow cytometry showed a greater than 10-fold reduction in the number of A4^ko^ T cells compared to WT T cells (Fig. 5I).

**Figure 5:**
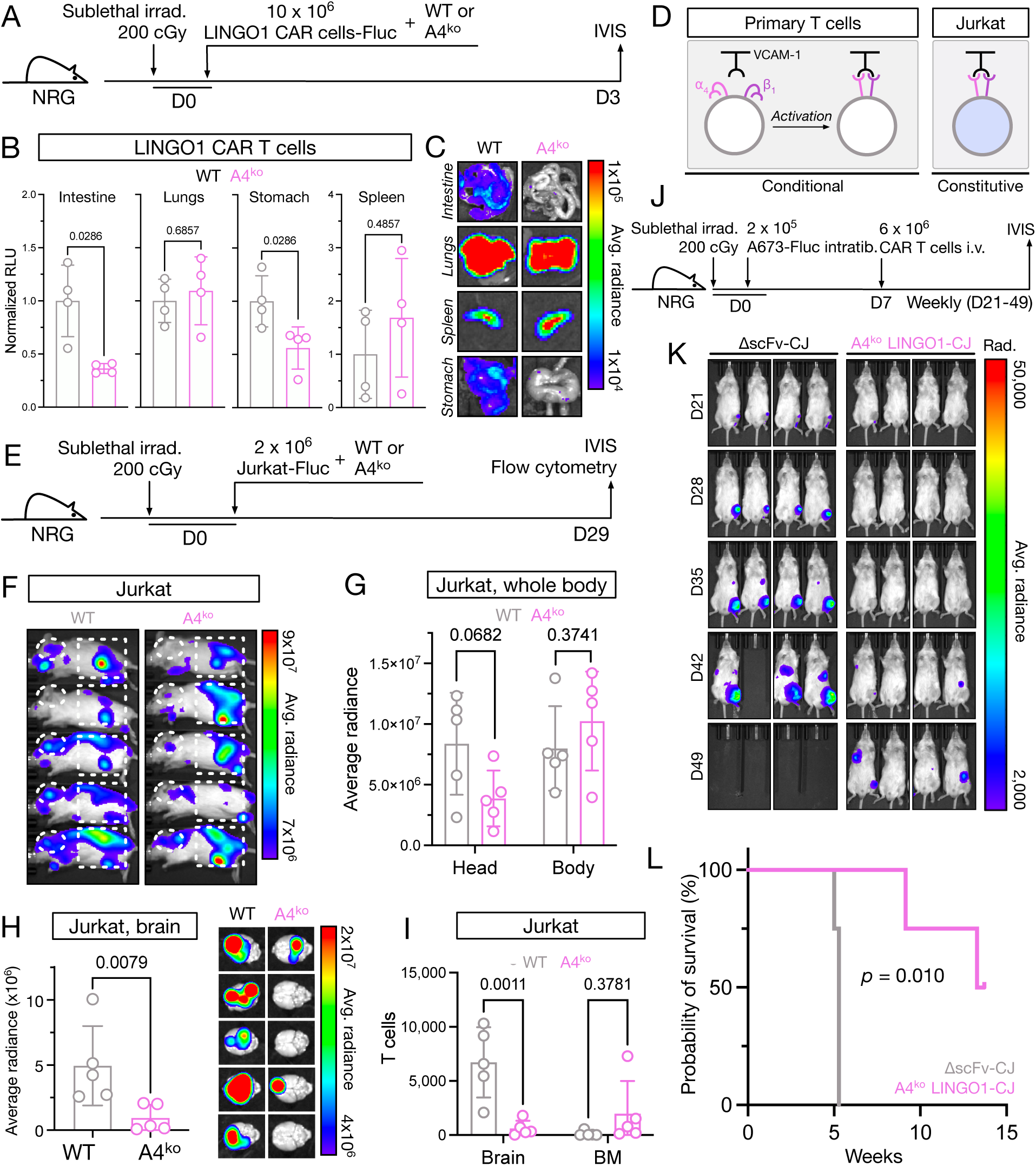
Integrin ⍺_4_ knockout prevents trafficking of CAR T cells to the CNS but not to ES tumors. **(A)** Schema of in vivo experiment to determine tissue trafficking of primary human T cells expressing firefly luciferase (Fluc). **(B)** Quantification of relative luminescence units (RLU) and representative example images of organs from one animal of explanted organs normalized to the mean signal of WT LINGO1 CAR T cells. Data represent mean ± S.D. from 4 animals. Statistical significance was determined by Mann-Whitney U test. **(C)** Example of luminescence observed in tissues from mice injected with A4^ko^ T cells. **(D)** Schema of integrin ⍺_4_β_1_ activation status in Jurkat and primary T cells. **(E)** Schema of in vivo experiment to determine CNS trafficking of A4^ko^ Jurkat cells. **(F)** In vivo imaging of mice injected with A4^ko^ Jurkat cells. Dotted lines indicate head and lower body regions of interest (ROI) used for luminescence quantification. **(G)** Quantification of luminescence signal of indicated ROIs from whole body imaging of mice injected with A4^ko^ Jurkat cells. Data represent mean ± S.D. from 5 animals per group. Statistical significance was determined by two-sided Student’s t test. **(H)** Quantification and images of luminescence signal measured in explanted brains from mice injected with WT of A4^ko^ Jurkat cells. Data represent mean ± S.D. from 5 animals per group. Statistical significance was determined by Mann-Whitney U test. **(I)** Number of T cells isolated from brains of mice injected with WT or A4^ko^ Jurkat cells. Data represent mean ± S.D. from 5 animals per group. Statistical significance was determined by two-way ANOVA. **(J)** Schema of orthotopic *in vivo* experiment to determine efficacy of A4^ko^ LINGO1 CAR T cells against ES. **(K)** Tumor burden in 4 mice per group bearing intratibial A673 tumors and treated with intravenous A4^ko^ LINGO1 CAR T cells or ΔscFv control CAR T cells as determined by IVIS. **(L)** Survival of ES tumor-bearing mice treated with LINGO1 CAR T cells. Statistical significance was determined by log-rank test.

Due to the absence of increased rates of cancer in patients treated with integrin ⍺_4_-targeting antibodies (*34*) we hypothesized that A4^ko^ may not be necessary for the trafficking of T cells to tumors located outside of the CNS. We therefore next investigated the efficacy of A4^ko^ LINGO1 CAR T cells against ES tumors *in vivo*. We had observed that systemically injected A673 cells frequently form metastases in the intestine (Suppl. Fig. S5D), in contrast to primary human ES tumors, which almost exclusively metastasize to the lungs or bone marrow (*30*). We therefore performed an orthotopic mouse model by delivering ES cells intratibially and CAR T cells systemically (Fig. 5J). A single dose of A4^ko^ LINGO1 CAR T cells was able to efficiently control ES tumors (Fig. 5K) and to significantly prolong survival (Fig. 5L), demonstrating that changes in CAR T cell adhesion behavior had selectively disrupted trafficking to defined normal tissues but not to ES tumors.

## DISCUSSION

Only two antigens, CD19 and BCMA, are currently being targeted by FDA-approved CAR T cell products and the list of promising candidate antigens, especially for solid tumors, is relatively short (*51–53*). This is in large part due to the ability of CAR T cells to kill cells expressing even low amounts of antigen (*9*), excluding any antigens from consideration that are also expressed in essential normal tissues. We demonstrate that the safety profile of CAR T cells targeting such shared antigens can be improved by altering their adhesion behavior, resulting in their efficient exclusion from defined healthy tissues.

In this work, we focused on the disruption of one of the most widely studied adhesion events, the interaction between integrin ⍺_4_β_1_ and VCAM-1. We demonstrate that A4^ko^ T cells are specifically prevented from trafficking to the CNS without altering tumor homing. To test the feasibility of this approach, we developed CAR T cells targeting LINGO1, an antigen expressed in ES and multiple cell types in the CNS of humans and mice. It has been shown that human integrin ⍺_4_β_1_ efficiently binds to mouse VCAM-1 (*43*) and we observed efficient integrin ⍺_4_-dependent trafficking of Jurkat T cells to mouse brains. However, in our *in vivo* studies we were unable to detect WT LINGO1 CAR T cells in the brains of mice or signs of overt neurotoxicity, even in our systemic ES model using WT LINGO1 CAR T cells. In addition to the likely limited steady-state trafficking of primary T cells to the CNS, targeting of LINGO1 may not result in overt neurotoxicity - a clinical trial of the LINGO1-targeting antibody opicinumab did not show an increased rate of serious adverse events compared to placebo (*27*) and LINGO1 knockout mice do not exhibit signs of abnormal development, locomotion, or behavior (*54*). We therefore determined the effect of A4^ko^ on CNS trafficking of T cells by developing a surrogate approach using the T cell line Jurkat, which expresses a constitutively active ⍺_4_β_1_ heterodimer (*50*) and has been shown to readily migrate to the CNS (*55*). In this experiment we observed a substantial reduction in CNS trafficking of T cells following A4^ko^. However, the extent of the effect of integrin ⍺_4_ knockout on CNS trafficking may differ in primary CAR T cells. Additional preclinical work demonstrating the safety of this approach, e.g. using non-human primates, will likely be necessary.

Analyzing the trafficking of LINGO1 CAR T cells to other normal tissues, we observed that A4^ko^ also prevented homing to the gastrointestinal tract, likely by disrupting formation of the integrin ⍺_4_β_7_ heterodimer. The primary ligand for integrin ⍺_4_β_7_ is mucosal vascular addressin cell adhesion molecule-1 (MAdCAM-1), which controls extravasation into mucosal tissues (*48*). An approach like the one presented in this work but specifically targeting integrin β_7_ could be used to selectively disrupt gastrointestinal trafficking to enable targeting of tumor antigens showing expression in the normal gastrointestinal tract.

In conclusion, we show that altering integrin composition and the resulting adhesion behavior of CAR T cells can be used to selectively restrict tissue homing. The approach allows the rational selection of antigens showing shared expression on normal tissues, and thereby expands the set of antigens targetable by CAR T cells.

## ONLINE METHODS

### Study approval

All recombinant DNA and biosafety work was approved by the institutional biosafety committees at University of Maryland Baltimore (protocol IBC-6040) and the University of Utah (protocol 37-14). Animal experiments were approved by the institutional animal care and use committee at the University of Utah (protocol 18-1104).

### Cell lines and primary human cells

A673, Jurkat, K562, SKES1, RDES, and Phoenix-Ampho cells were purchased from ATCC and cultured according to ATCC instructions. Lenti-X 293T cells were purchased from Takara and cultured according to the manufacturer’s instructions. Cell lines were authenticated by their respective supplier. Healthy donor buffy coats were obtained from the Blood Centers of America and the New York Blood Center. Peripheral blood mononuclear cells were isolated from buffy coats by density gradient using FicollPaque (GE) as previously described (*19*).

### Production of transgenic cell lines

For cytotoxicity, adhesion, and trafficking studies, A673, Jurkat, SKES1, and RDES cell lines were transduced with firefly luciferase (Fluc) as previously described (*56*) using lentivirus based on pHIV-Luc-ZsGreen or a variant of this plasmid in which zsGreen was replaced with mCherry and cells were sorted on a 4-laser FACSAria II cell sorter (BD) for GFP expression. K562-Fluc cells were also transduced to express full-length human (accession # NP_116197.4) or mouse (accession # NP_001348044.1) LINGO1 and EGFP with SFG-based gammaretroviral expression constructs and mCherry^+^EGFP^+^ cells were sorted on a FACSaria flow cytometer.

To achieve integrin knockout in Jurkat-Fluc cells, a lentiviral transfer plasmid was constructed expressing hSpCas9 and mTag-BFP2 separated by a P2A sequence under control of the EFS-NS promoter and an individual guide RNA (gRNA) under control of the human U6 promoter. The following target sequences were used to generate gRNAs: <u>ITGA4.2</u>: 5’-CAG CAT ACT ACC GAA GTA GT-3’; <u>ITGA4.3</u>: 5’-GTG TTT GTG TAC ATC AAC TC-3’; <u>ITGB1.1</u>: 5’-TGC TGT GTG TTT GCT CAA AC-3’; <u>ITGB1.5</u>: 5’-TTT GCT GGA GAT GGG AAA CT-3’. A gRNA without a targeting sequence was used as the non-specific (NS) control. Jurkat cells were transduced with these constructs and sorted for the presence of BFP and the absence of integrin ⍺_4_ or β_1_ on a FACSaria flow cytometer.

### Identification of single-tissue surface antigens and Ewing sarcoma RNA expression

Consensus tissue gene expression values combining RNA sequencing data from the Human Protein Atlas and the Genotype Tissue Expression projects were obtained from the Human Protein Atlas website (proteinatlas.org) on https://www.proteinatlas.org/download/rna_tissue_consensus.tsv.zip (created: 17 June 2023). The multiple CNS, hematopoietic, and intestinal tissues present in this dataset were collapsed into single “CNS”, “hematopoietic”, and “intestine” tissues using the highest expression value within the respective group. All antigens showing an expression of greater than 50 normalized transcripts per million (nTPM) in one tissue and expression of less than 20 nTPM in all other tissues were selected as single-tissue antigens.

Single-tissue surface antigens (STSAs) were identified by mapping gene symbols of single-tissue antigens to UniProt KB identifiers and selecting genes with evidence of protein expression, presence of a transmembrane domain, and containing the term “cell membrane” in the subcellular localization field in UniProt (accessed on 9 December 2023). Duplicate entries returned by UniProt were removed manually. The resulting list was used to create a matrix of STSAs and cancer gene expression using RNA HPA cell line cancer gene data available from the Human Protein Atlas website on https://www.proteinatlas.org/download/rna_celline_cancer.tsv.zip (created: 17 June 2023).

RNA expression of LINGO1 in osteosarcoma and Ewing sarcoma (ES) cell lines was obtained from the TRON Cell Line Portal (TCLP) dataset available on https://github.com/TRON-Bioinformatics/TCLP (accessed on 28 August 2023) and validated by RT-PCR.

RNA expression of LINGO1 in normal tissues was obtained through the NCBI Gene portal for human (Gene ID: 84894) and mouse (Gene ID: 235402) tissues. Brain cell type-specific expression data was obtained from the Human Protein Atlas website on https://www.proteinatlas.org/ENSG00000169783-LINGO1/brain (accessed on 19 April 2023).

RNA expression data of LINGO1 in archived human ES tumor biopsies, normal tissues, and neural crest and mesenchymal stem cells was obtained from the R2 Genomics Analysis and Visualization Platform on https://hgserver1.amc.nl/ by accessing Affymetrix Exon array data available through the NCBI Gene Expression Omnibus (GEO) platform (accession # GSE68776 (*20*)).

Single-cell RNA expression of LINGO1 and other ES-associated antigens was obtained through the NCBI GEO platform (accession # GSE130024 (*25*)). The dataset was processed, explored, and visualized using Cellenics® community instance (https://scp.biomage.net/) hosted by Biomage (https://biomage.net/).

### Single-chain variable fragment (scFv) expression and purification

ScFvs based on LINGO1-specific monoclonal antibodies Li62 and Li81 synthesized by GeneArt (Thermo Fisher) and cloned into pSANG10. ScFvs were expressed overnight in 25 mL MagicMedia E. coli autoexpression medium (Thermo-Fisher). Periplasmic extracts were generated from autoinduction cultures as previously described (*57*). ScFvs were purified by immobilized metal affinity chromatography using NiNTA resin (Thermo-Fisher). Concentrations of purified scFvs were determined by bicinchoninic acid (BCA) protein assay (Thermo-Fisher). Purity of scFvs was confirmed by SDS-PAGE and GelCode^TM^ Blue staining (Thermo Fisher).

### Flow cytometry expression analysis

Flow cytometry staining and analyses were performed as previously described (*56*, *57*). Indirect staining was used for the analysis of scFv binding to ES cell lines by using a fluorophore-conjugated secondary anti-FLAG tag antibody. Antibodies used for flow cytometry analyses are listed in Supplementary Table S3. Commercially available antibodies were used at dilutions recommended by the respective manufacturer. For the analysis of phenotypic, activation, and exhaustion markers on A4^ko^ CAR T cells, data was acquired on a full spectrum Aurora flow cytometer (Cytek). All other flow cytometry data was acquired on an LSR Fortessa or LSR II flow cytometer (BD) and all data was analyzed using FlowJo 10 (BD).

### Quantification of T cells in mouse brain tissue

Explanted brains from animals injected with A4^ko^ or WT Jurkat-Fluc cells were washed twice in PBS and subsequently homogenized using, first, a 70 μm cell strainer followed by a 18G syringe. Lymphocytes were isolated using a 70% Percoll gradient. Isolated lymphocytes were stained using antibodies for huCD45, msCD45, and huCD3 and DAPI for live/dead gating. The number of DAPI^-^msCD45^-^huCD45^+^huCD3^+^ T cells per 1,000 AccuCheck counting beads (Thermo Fisher) was quantified by flow cytometry using an LSR II flow cytometer (BD).

### Interferon gamma (IFN-γ) Enzyme-linked Immunosorbent Assay (ELISA)

Supernatants were harvested from 96-well 16-hour co-cultures and immediately frozen at −80°C. IFN-γ concentrations were determined by standard curve using a commercial ELISA kit according to the manufacturer’s instructions (Biolegend). Absorbance was measured on a multi-mode plate reader (Tecan).

### Western blot

Total lysates were extracted from CAR T cells on day 9 of production using RIPA buffer (Thermo Fisher). Protein concentrations were determined by BCA assay (Thermo-Fisher). Samples were separated by sodium dodecyl sulfate–polyacrylamide gel electrophoresis, and separated proteins transferred to nitrocellulose membranes using an iBlot2 transfer system (Thermo Fisher). Membranes were blocked with 5% non-fat milk-TBS and incubated with primary antibodies against BATF3, pc-Jun (S73), c-Jun, and β-actin (see Supplementary Table S3). Membranes were washed and developed using species-specific secondary anti-IgG/HRP antibodies (R&D Systems) and Western Lightning Plus-ECL solution (PerkinElmer). Bands were visualized on an iBright 1500 imaging system (Thermo).

### Luciferase-based cytotoxicity assay

For cytotoxicity assays, 4-5×10^4^ target cells were seeded in each well of a round bottom (K562) or black, flat bottom (A673, SKES1, RDES) 96-well plate. Indicated ratios of CAR T cells were co-cultured with target cells for 16 hours at 37°C/5% CO_2_. After the co-culture, cells were suspended by gentle pipetting and 100 μL were moved to a 96-well black flat bottom plate. 100 μL D-luciferin at 150 μg/ml (Gold Biotechnology, cat. no. LUCNA-2G) was added to the cells and incubated for 5 minutes at 37°C. Luminescence was determined on a Spark multi-mode plate reader (Tecan).

### CAR T cell production

ScFvs were cloned into a previously described second generation 4-1BB-based CAR construct (*19*) in the gammaretroviral SFG backbone or the lentiviral pRRL backbone. For some constructs, the full-length sequence of human c-Jun (Uniprot P05412) was synthesized and cloned upstream of the respective CAR separated by a P2A sequence (Twist Bioscience). Lentivirus or amphotropic gammaretrovirus was generated by transfection of Lenti-X 293T cells or Phoenix-Ampho cells (ATCC cat. No. CRL-3213) using the respective transfer plasmids and Lipofectamine 2000 according to the manufacturer’s instructions. Virus-containing supernatants were concentrated with Lenti-X or Retro-X concentrator (Takara), respectively. PBMCs were stimulated for 2 days with CD3/CD28 T Activator beads (Thermo-Fisher, cat. no. 11131D) in the presence of 40IU/ml IL-2 (R&D Systems, cat. no. 202-IL-010) in AIM V media (Thermo-Fisher) supplemented with 5% human serum (Sigma, cat. no. H3667) and incubated at 37°C/5% CO_2_. Bead-stimulated cells were transferred to Retronectin-coated (Takara) virus-containing plates and incubated overnight. Transduction was repeated the next day before counting and diluting cells to 0.4 x 10^6^ cells/ml. To generate CAR T cells expressing Fluc for trafficking studies, CAR gammaretrovirus and pHIV-Luc-zsGreen-based lentivirus were immobilized on Retronectin simultaneously at a v/v ratio of 1:9 (gammaretrovirus:lentivirus).

To generate A4^ko^ CAR T cells, on day 4, anti-CD3/CD28 T Activator beads (Thermo Fisher) were removed from the transduced cells by magnet. CRISPR RNP particles containing 25 μg Cas9 only (mock/WT) or Cas9 and 150 pmol synthetic ITGA4-specific TrueGuide gRNA (target sequence: 5’-GTGTTTGTGTACATCAACTC-3’) were prepared according to the manufacturer’s instructions (Thermo). 3 x 10^6^ cells were electroporated in the presence of CRISPR-RNPs using the NEON electroporation system (Thermo) using the following settings: 1,600 V, 10 ms, 3 pulses. Following the electroporation, cells were transferred to prewarmed media and incubated for 2 hours at 37°C. Following the incubation, new anti-CD3/CD28 T cell activation beads were added to the electroporated cells at a ratio of 1:1, and cells were grown for an additional 6 days before removing beads using a DynaMag-15 magnet (Thermo Fisher). IL-2 was replenished every 2 days to 40 IU/ml. Cells were frozen in 90% FCS/10% DMSO and stored in liquid nitrogen.

### In vivo tumor models

Six- to 8-week-old male NOD.Cg-Rag1^tm1Mom^ Il2rg^tm1Wjl^/SzJ (NRG, The Jackson Laboratory) mice were irradiated with a sublethal dose of 200 cGy (Rad-Source RS-2000) and injected on the same day with the indicated numbers of A673-Fluc cells. For the metastatic ES model tumor cells were injected through the lateral tail vein. For the orthotopic model, tumor cells were injected intratibially. On day 7 after tumor cell injection, the indicated number of LINGO1 CAR T cells or CAR T cells lacking a binding domain (ΔscFv) were injected into the lateral tail vein. To determine CAR T cell efficacy, animals were weighed twice weekly and monitored for signs of distress in accordance with institutional regulations. For *in vivo* imagining, mice received an intraperitoneal injection of 3.3 mg D-luciferin (GOLDBIO # LUCK-10G). Photographic and luminescent images were acquired starting 10 minutes after the D-luciferin injection, both in prone and supine position using an *in vivo* imaging system (IVIS). ES progression was monitored every 7 days until the study endpoint. Average radiance (p/s/cm²/sr) was quantified for individual animals using Living Image 4.8 software (PerkinElmer).

### VCAM-1 adhesion assay

To determine binding of T cells to VCAM-1 following integrin knockout, black 96-well flat bottom MaxiSorp plates (Nunc) were coated with 50 μl of 5 μg/ml recombinant VCAM-1-Fc (R&D Systems, cat # 862-VC) over night at 4C. The next day, wells were washed and blocked with 5% human AB serum in PBS for 1 hour at room temperature. Wells were washed twice with PBS and 1×10^5^ Jurkat-Fluc cells were added to the wells. Plates were incubated for 4 hours in complete media at 37C/5% CO_2_. Following incubation, wells were washed the indicated number of times using an automated plate washer (Tecan HydroFlex) with 300 μl PBS at 200 μl/s. The remaining Jurkat-Fluc cells per well were quantified by luminescence as described for the luciferase-based cytotoxicity assay using an EnVision plate reader (PerkinElmer). Values were normalized to the “no wash” condition.

### CodePlex secretome assay

Supernatants were harvested from overnight co-cultures and stored at −80°C until further use. On the day of the assay, samples were thawed and 11 μl of supernatant per sample was added to CodePlex Human Adaptive Immune secretome chips (Isoplexis). Chips were loaded into the Isolight reader and cytokines measured using default settings. Automated analysis of raw data was performed using IsoSpeak (Isoplexis).

### Reverse transcriptase polymerase chain reaction (RT-PCR)

LINGO1 expression was measured by reverse transcription polymerase chain reaction (RT-PCR). RNA from ES cell lines was isolated using the RNeasy Mini kit (Qiagen), according to the manufacturer’s instructions. Total RNA from normal human brain tissue was obtained commercially (Thermo Fisher). Complementary DNA (cDNA) was generated using the SuperScript III First-Strand Synthesis system (Thermo). We performed PCRs for LINGO1 (<u>LINGO1_F</u>: 5ʹ-CCC GAG ATG CAG GTG AGC-3ʹ; <u>LINGO1_R</u>: 5ʹ-CTA GCG GGA TGA GCT TCA GG-3ʹ) and GAPDH (<u>GAPDH_F</u>: 5ʹ-ATT GCC CTC AAC GAC CAC TTT G-3ʹ; <u>GAPDH_R</u>: 5ʹ-TTG ATG GTA CAT GAC AAG GTG CGG-3ʹ) using MyTaq^TM^ Red Mix (Bioline) and a ProFlex thermal cycler PCR system (Thermo). We performed qualitative gel electrophoresis using SYBR Safe DNA stain and imaged gels using a Gel Doc XR+ System (Bio-Rad).

### Bead-based depletion of residual integrin ⍺_4_+ LINGO1 CAR T cells

A4^ko^ LINGO1 CAR T cells were washed with 5% BSA/PBS and incubated with a biotinylated anti-integrin ⍺_4_ antibody (1 μg/1×10^6^ cells) for 30 minutes on ice. Cells were washed with 5% BSA/PBS and transferred to a microcentrifuge tube. Streptavidin Dynabeads (Thermo, cat. no. 65305) were added to the cells (10 μl/1 x 10^6^ cells), vortexed, and incubated for 20 minutes on an end-over-end shaker at 4C. Next, the tube was placed on magnet for 2 minutes and supernatant containing the purified cell population was transferred to a new tube before proceeding to downstream assays.

### In vivo trafficking experiments

To determine trafficking of WT and A4^ko^ LINGO1 CAR T cells expressing Fluc, NOD.*Cg-Rag1^tm1Mom^Il2rg^tm1Wjl^*/SzJ (NRG) mice were sublethally irradiated and injected with the indicated number of CAR T cells. After 3 days, intestine, spleen, brain, stomach, and lungs were explanted and imaged by IVIS.

To determine CNS trafficking of T cells, NRG mice were sublethally irradiated and injected with 2 x 10^6^ WT or A4^ko^ Jurkat-Fluc cells by tail vein. Animals were imaged by IVIS on day 29 and subsequently euthanized. Brains were explanted and imaged individually by IVIS. Radiance values were determined using LivingImage 4.8 software (PerkinElmer).

### Statistical analysis and data visualization

The respective statistical tests are stated in the figure legends. Generally, the significance of differences of continuous variables between two groups was determined by two-sided Student’s t test or Mann-Whitney U test. Significance of differences between more than two groups were determined by one- or two-way analysis of variance (ANOVA). Differences in luminescence at individual time points during *in vivo* experiments were determined by Mann-Whitney U test. Statistical significance of differences in survival was determined by log-rank test. All statistical tests were performed using Prism 10 (GraphPad Software). Results were considered significant when *p* or adjusted *p* < 0.05.

## ACKNOWLEDGEMENTS

This publication was supported by funds through the Maryland Department of Health’s Cigarette Restitution Fund Program (CH-649-CRF) and the National Cancer Institute Cancer Center Support Grant (P30CA134274). This publication was supported by an NIAID-funded predoctoral fellowship to JMB (T32 AI095190). We thank the Huntsman Cancer Institute at the University of Utah for the use of the Preclinical Research Resource (PRR) and the Flow Cytometry core facility. We also thank the University of Maryland School of Medicine & Greenebaum Comprehensive Cancer Center’s Flow Cytometry core, which assisted with flow cytometry analyses and cell sorting. The plasmid pHIV-Luc-ZsGreen was a gift from Dr. B. Welm (Addgene #39196). The plasmid SFG.CNb30_opt.IRES.eGFP was a gift from Dr. M. Pule (Addgene #22493). The plasmid pSANG10-3F (Addgene plasmid #39264) was a gift from Dr. J. McCafferty. The graphical abstract was generated in part using BioRender.com.

## AUTHOR CONTRIBUTIONS

EAM and TL conceived the project. EAM generated CAR constructs and LINGO1 CAR T cells, determined LINGO1 RNA and surface expression in ES cell lines, engineered lentiviral integrin-targeting CRISPR/Cas9 constructs, determined LINGO1 CAR surface expression and *in vitro* cytotoxicity, and performed all Jurkat-based integrin ⍺_4_ in vitro assays including flow cytometry and VCAM-1 adhesion assays. KAD generated LINGO1 CAR T cells, performed flow cytometry analysis, developed the *integrin ⍺-4* depletion method, and analyzed data. JMB performed cytotoxicity assays, analyzed data, and revised the manuscript. TI performed the CodePlex assay and analyzed data. WS and ER planned and performed in vivo tumor models and trafficking experiments, and analyzed data. XF developed the full-spectrum flow cytometry panel and performed the flow cytometry analysis. SVA, FI, SVR, AW, EVM, and MLO planned experiments and analyzed data. DA planned and supervised CodePlex cytokine analyses and analyzed data. TL supervised all work related to this project, performed computational analyses, generated A4^ko^ and c-Jun-overexpressing LINGO1 CAR T cells, performed flow cytometry assays and western blots, generated LINGO1-expressing K562 T cells, quantified T cells in mouse brain tissues, and wrote the manuscript. All authors reviewed and approved the manuscript.

## DATA AVAILABILITY

All original data associated with this study are in the paper or supplementary materials. Publicly available gene expression data obtained from the Human Protein Atlas project, the TRON Cell Line Portal, and the NCBI GEO database are available from the respective websites. The plasmids, including LINGO1 CAR expression and lentiviral integrin ⍺_4_ CRISPR/Cas9 constructs, generated as part of this study are available under a Material Transfer Agreement from the corresponding author.

