## Supplementary Material for "Restricting CAR T Cell Trafficking Expands Targetable Antigen Space"

Morales et al.

**SUPPLEMENTARY FIGURES**

**Supplementary Figure S1: Development of CAR T cells targeting the CNS-restricted STSA LINGO1. (A)** Average expression of STSAs per normal tissue in human cancer cell lines as determined by RNAseq. nTPM: normalized transcripts per million. Grey bars indicate cancer types primarily located within the respective normal tissue. **(B)** mRNA expression of *LINGO1* and *GAPDH* housekeeping gene in Ewing sarcoma cell lines and healthy brain tissue as determined by RT-PCR. **(C)** Single-cell RNA sequencing of 4 patient-derived ES xenograft samples (*25*) indicating intratumoral expression of conventional ES markers EWSR1, FLI1, and CD99 as well as 10 candidate CAR T cell antigens. **(D)** Concentration of NiNTA-purified LINGO1-specific scFvs as determined by BCA protein assay. Data represent mean ± S.D. from 2 technical replicates. **(E)** Surface LINGO1 expression in ES cell line A673 stained with opicinumab or Li81-LH scFv and a respective APC-conjugated secondary antibody as determined by flow cytometry. **(F)** LINGO1 CAR surface expression on primary human T cells transduced with lentiviral expression constructs after staining with an anti-HA/APC antibody as determined by flow cytometry. **(G)** Schema of gammaretroviral construct used for the simultaneous expression of a LINGO1 or ∆scFv CAR and c-Jun. **(H)** Surface expression of LINGO1 in parental K562 cells, or K562 cells transduced with a human LINGO1 (huLINGO1) or mouse LINGO1 (msLINGO1) expression construct as determined by flow cytometry. **(I)** Homology of human (UniProt Q96FE5) and mouse LINGO1 (UniProt Q9D1T0) as determined by blastp.

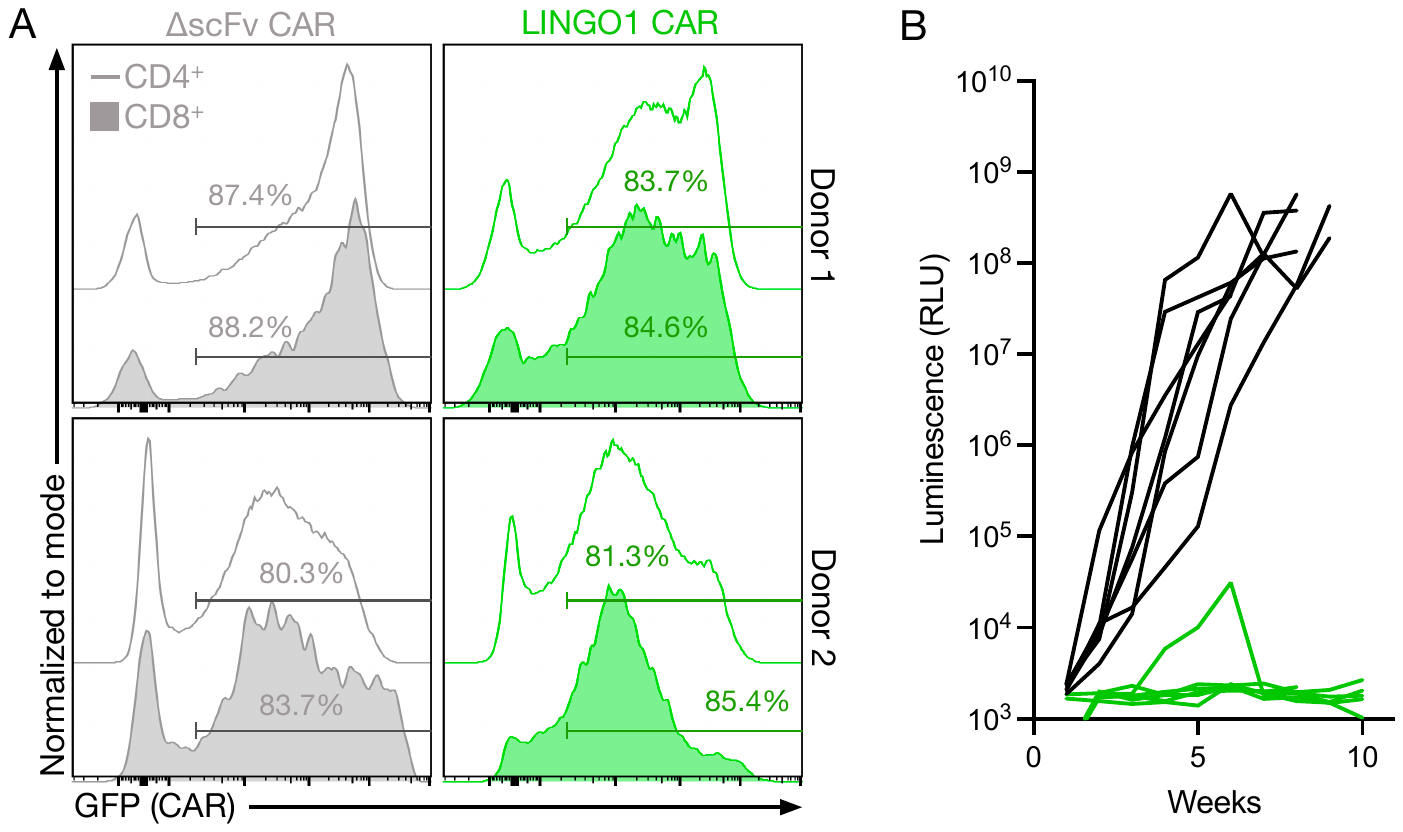

**Supplementary Figure S2: LINGO1 CAR T cells are effective against Ewing sarcoma. (A)** LINGO1-CJ CAR transduction efficiency of healthy human donor CD4^+^ or CD8^+^ T cells by gammaretrovirus by the end of CAR T cell manufacturing as determined by full-spectrum flow cytometry. **(B)** Tumor burden of mice systemically injected with A673-Fluc cells and LINGO1-CJ (*n* = 6) or ∆scFv (*n* = 7) CAR T cells as determined by in vivo imaging system.

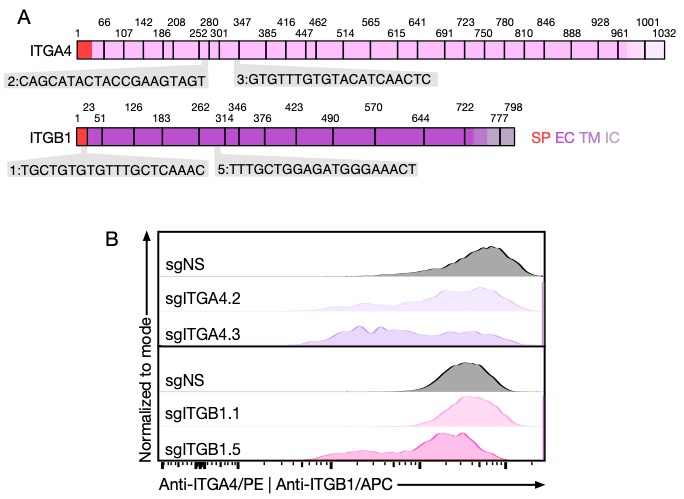

**Supplementary Figure S3: CRISPR/Cas9-mediated integrin ⍺_4_β_1_ knockout in T cells. (A)** Exon structure of integrin ⍺_4_ and integrin β_1_ and respective gRNA sequences. SP signal peptide; EC extracellular domain; TM transmembrane domain; IC intracellular domain. **(B)** Integrin ⍺_4_ and integrin β_1_ expression on the surface of Jurkat T cells after lentiviral transduction with Cas9 and individual gRNAs targeting *ITGA4* (sgITGA4), *ITGAB1* (sgITGB1), or a non-specific negative control gRNA (sgNS) as determined by flow cytometry.

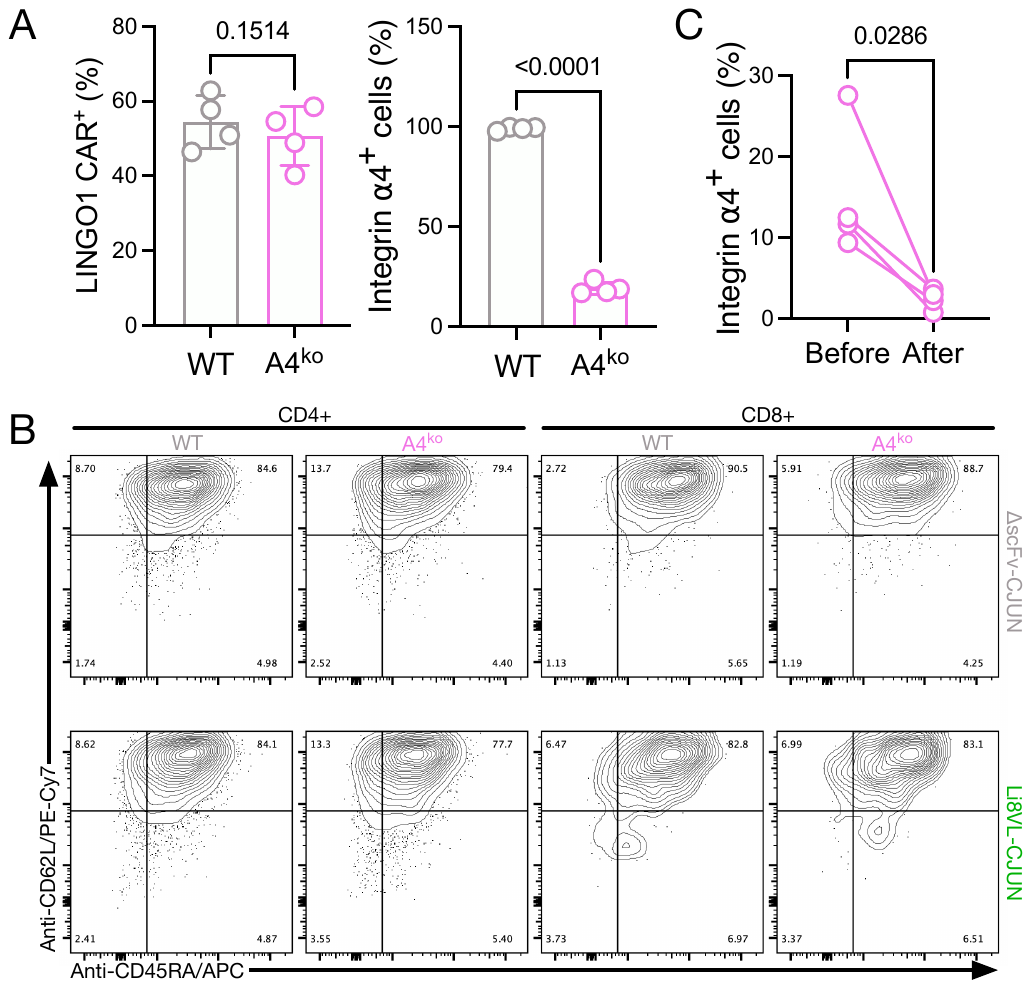

**Supplementary Figure S4: Effect of integrin ⍺_4_ knockout on CAR T cell phenotype. (A)** Percentage of LINGO1 CAR^+^ or integrin ⍺_4_^pos^ cells in CAR T cell products at the end of manufacturing as determined by flow cytometry. Data represent mean ± S.D. from 4 separately manufactured CAR T cell products. Statistical significance was determined by paired two-tailed *t* test. **(B)** Phenotype of CAR T cell products at the end of manufacturing as determined by flow cytometry. **(C)** Efficiency of integrin ⍺_4_ bead depletion as determined by quantifying integrin ⍺_4_^pos^ T cells before and after depletion. Data represent mean ± S.D. from 4 separately manufactured CAR T cell products. Statistical significance was determined by Mann-Whitney U test.

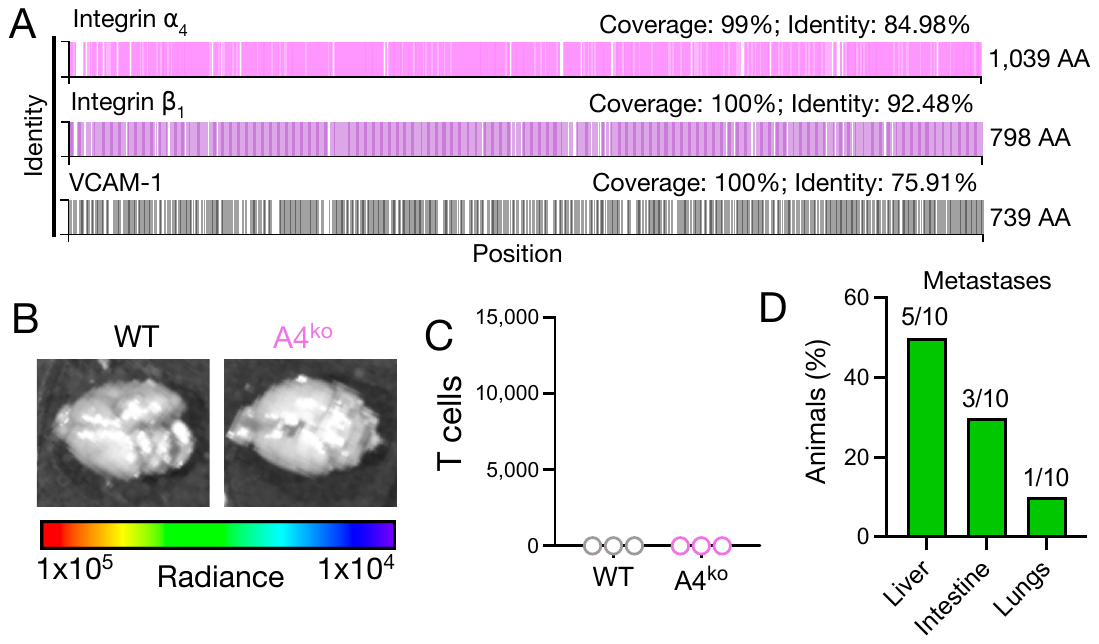

**Supplementary Figure S5: In vivo trafficking of A4^ko^ T cells. (A)** Homology between human and mouse integrin ⍺_4_ (UniProt P13612 and Q00651), integrin β_1_ (UniProt P05556 and P09055), and VCAM-1 (UniProt P19320 and P29533) as determined by blastp. **(B)** Luminescence as determined by IVIS in 1 animal per condition and **(C)** numbers of T cells as determined by flow cytometry in explanted brains from 3 animals per group injected with WT or A4^ko^ primary human LINGO1 CAR T cells expressing Fluc. Data represent mean ± S.D. from 3 animals per group. **(D)** Percentage of organs macroscopically showing tumor metastases in mice intravenously injected with A673-Fluc cells and ∆scFv-CJ CAR T cells. Numbers above bars indicate numbers of animals with metastases in the respective organ/total number of animals.

**SUPPLEMENTARY TABLES**

**Supplementary Table S1: List of single-tissue antigens.**

**Supplementary Table S2: Matrix of single-tissue surface antigens and their expression in cancer.**

| **Target** | **Clone** | **Fluorophore** | **Supplier** | **Target cells** |
| --- | --- | --- | --- | --- |
| LINGO1 | Opicinumab | None | ProSci | Ewing |
| Integrin ⍺_4_ | 9F10 | APC/PE/Biotin | Biolegend | CAR T |
| Integrin β_1_ | TS2/16 | APC | Biolegend | CAR T |
| CD3 | UCHT1 | BV711/BV421/  BUV496 | Biolegend | CAR T |
| huCD45 | 2D1 | APC | Biolegend | CAR T |
| msCD45 | 30-F11 | APC/PE-Cy7 | Biolegend | CAR T |
| CD45RA | HI100 | APC | Biolegend | CAR T |
| CD62L | DREG-56 | PE-Cy7 | Biolegend | CAR T |
| CD3 | SK7 | Spark 550 | Biolegend | CAR T |
| CD4 | SK3 | BV805 | BD | CAR T |
| CD8 | RPA-T8 | BV510 | Biolegend | CAR T |
| CD27 | O323 | APC/Fire750 | Biolegend | CAR T |
| CD45RA | HI100 | PerCP | Biolegend | CAR T |
| CD45RO | UCHL1 | BV750 | Biolegend | CAR T |
| CD56 | NCAM16.2 | BUV563 | BD | CAR T |
| CD62L | SK11 | BUV395 | BD | CAR T |
| CD95 | DX2 | Alexa Fluor 700 | Biolegend | CAR T |
| CD197/CCR7 | G03H7 | BV650 | Biolegend | CAR T |
| CD25 | BC96 | PE-Cy5 | Biolegend | CAR T |
| CD38 | HB7 | BUV737 | BD | CAR T |
| CD127 | A019D5 | BV421 | Biolegend | CAR T |
| LAG-3 | 11C3C65 | PE-Cy7 | Biolegend | CAR T |
| TIM-3 | F382E2 | BV605 | Biolegend | CAR T |
| PD-1 | EH12.2H7 | APC | Biolegend | CAR T |
| Hemagglutinin tag | 6E2 | PE/APC | Cell Signaling Technology | CAR T |
| BATF3 | Polyclonal  # AF7437 | Unconjugated | R&D Systems | Western blot |
| c-Jun | 60A8 | Unconjugated | Cell Signaling Technology | Western blot |
| pc-Jun (S73) | D47G9 | Unconjugated | Cell Signaling Technology | Western blot |
| β-actin | 937215 | Unconjugated | R&D Systems | Western blot |
| 7-AAD |  | N/A | Biolegend | Live/dead |
| DAPI | D1306 | N/A | Life Technologies | Live/dead |
| DYKDDDDK | M2 | PE | Sigma-Aldrich | Multiple |
| DYKDDDDK | L2 | PE/APC | Biolegend | Multiple |

**Supplementary Table S3: Table of monoclonal antibodies and viability dyes used for flow cytometry analyses.**
